# Detection and characterization of replication origins defined by DNA polymerase epsilon

**DOI:** 10.1101/2021.07.27.453931

**Authors:** Roman Jaksik, David A. Wheeler, Marek Kimmel

## Abstract

**Background:** Although the process of DNA replication is highly conserved the location of origins of replication (ORI) may vary from one tissue to the next or one round of replication to the next in eukaryotes, suggesting flexibility in the choice of locations to initiate replication. Lists of human ORI therefore vary widely in number and location and there are no methods available to compare them.

**Results:** Here we report the genome-wide localization of ORI in POLE-mutated human tumors using whole genome sequencing data. Mutations accumulated after many rounds of replication of unsynchronized dividing cell populations in tumors allow to identify constitutive origins, which we show are shared with high-fidelity between individuals and tumor types. Using a Smith–Waterman-like dynamic programming approach, we compared replication origin positions obtained from multiple different methods. The comparison allowed us to define a consensus set of replication origins, identified consistently by multiple ORI detection methods.

**Conclusions:** Many DNA features co-localized with the consensus set of ORI, including chromatin loop anchors, G-quadruplexes, S/MARs and CpGs. Among all features, the H2A.Z histone exhibited the most significant association. Our results show that mutation-based detection of replication origins is a viable approach to determining their location and associated sequence features.

## Background

DNA replication origins are crucial for initiation of the DNA synthesis, guiding the recruitment of proteins that form the pre-replication complex (pre-RC), including Mcm2-7 helicase. Helicase leads to the creation of a replication bubble, making the DNA accessible to polymerases, which replicate DNA in a bidirectional manner (3). The first step in pre-RC formation is recruitment of the origin recognition complex (ORC) that binds to specific regions in the DNA. These regions, referred to as the DNA replication origins (ORI) are selected based on sequence specificity in yeast (4, 5); however, in humans the recognition mechanism utilizes various DNA characteristics (7), and only a limited number of origins are active at each cell cycle (8). Efficiency of activation of the origins is used to classify them into three categories: constitutive, flexible and dormant (7). Constitutive origins are used by all cells, independent of the cell type in each cell cycle, whereas activation of the flexible origins may vary in position or from one cell cycle to the next, or one cell type to the next (7). Dormant origins become active in stress conditions that affect the S phase, including serum starvation and DNA damage (9). Variation in origin activation by Flexible or Dormant origins may be one reason for large differences in the number of ORI between different cell types (10-12) and could also help explain differences in results obtained using various methods. Large variation in numbers and positioning of ORI in higher organisms constitutes one of the major confounding factors in the study of human ORI.

The target recognition mechanism of ORC requires DNA characteristics similar to those required by transcription factors at transcriptions start sites (TSS). They include: nucleotide composition, chromatin state, DNA methylation, and secondary structure of DNA (7, 13). Since ORI must be accessible to protein binding, their locations were shown to coincide with the nucleosome free regions, histone acetylation and DNAse sensitive sites (14). Additionally, a low DNA methylation level is an important factor, making some promoter regions suitable targets for ORC binding (15). As a result. several of the best studied ORI are in the vicinity of TSS of known genes such as *MYC, TOP1* and *LMNB2* (16-18).

Despite the similarities to TSS there exists no definite evidence of the existence of specific sequence motifs required to initiate ORC assembly in humans. However, in some unicellular eukaryotic genomes *Cis*-acting sequences determine the location of replication origins (4) and in yeast two sequence elements are necessary: a 17-bp autonomously replicating consensus sequence (ACS) that binds Origin Recognition Complex (ORC): WWWWTTTAYRTTTWGTT (19) and a broader sequence context encompassing 200 to 300 bp that appears to be important for depleting nucleosomes from the origin (20-23). In human cells, nucleotide composition (2, 24, 25) and G-quadruplexes (26) have been shown to increase the replication origin activity also affecting its location. DNA structure is also believed to play an important role in replication origin location and activity. Chromatin loops were shown to be associated with origins (27), and also it is believed that DNA replication is initiated in regions attached to the nuclear matrix (MARs) (28). In a more recent work DNA replication origins were also shown to be overrepresented at the borders of topologically associating domains (TADs) (25).

Previously it was proposed (29) that mutational patterns emanated by the replicative DNA polymerases might effectively map the origins of replication, using mutations identified in whole genome sequencing experiments. Using this information, we developed a method for the detection of constitutive origins of replication, mORI (mutationally-defined ORI), based solely on mutation data from mutator phenotype tumor genomes. We also developed a novel method for the comparison of genomic positions which we used to compare multiple replication origin detection methods. Finally, we used the identified replication origins to characterize DNA structure in the vicinity of constitutive replication origins, determining the factors that are associated with their location.

## Results

### Detection of replication origins based on mutation patterns

DNA Polymerase ε is assumed to be responsible for the leading strand synthesis (30), a result first discovered in yeast, and subsequently reinforced by strong strand biases of context-specific mutations observed in cancers with mutation in the proof-reading domain of the enzyme – Fig. 1A (29, 31). To capture this feature and to identify potential replication origins we analyzed the distribution of mutations in POLE exonuclease damage tumors by developing a POLE-exo associated Mutation Asymmetry score (PMA). We obtained whole genome sequencing data (WGS) of tumors harboring POLE-exo mutation generated by the TCGA and ICGC projects (Supplementary Table 1) and focused on 20 cases out of 43 in which at least 20% of all mutations are either TCT→TAT and TCG→TTG — the two mutations most commonly found in these patients ((32) see mutation signatures 10a and 10b).

**Fig. 1:**
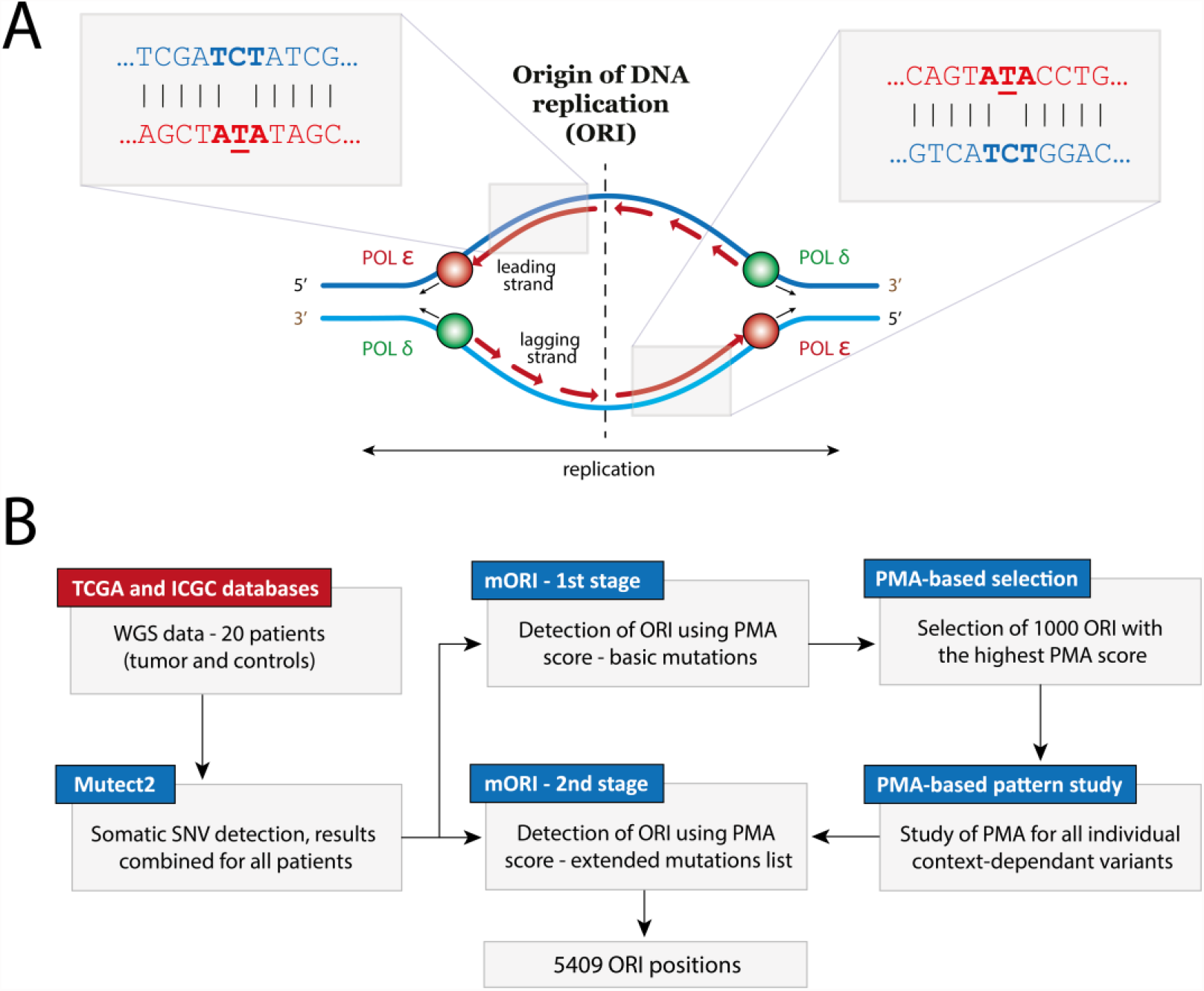
**A)** DNA replication in human cells. Polymerase ε synthesizes the leading strand while polymerase δ is responsible for the synthesis of the lagging strand (30). Both polymerases start the synthesis from the replication origin (ORI) after replication is initiated by the origin recognition complex (ORC). Damage in the exonuclease domain of polymerase ε lead to an increased mutation rate with a preference for C→A mutation in TxT context, observed as T<u>C</u>T->T<u>A</u>T the newly replicated strand. This creates an asymmetric pattern of TCT>TAT and AGA>ATA in the vicinity of replication origins. **B)** Diagram of the two-stage ORI detection method (mORI), based on the PMA score

Although TCT→TAT and TCG→TTG variants were reported to make up the majority of all mutations in POLE-exo mutant tumors, we noticed that other mutation types might be of relevance since they also occurred in similar spatial patterns around putative ORIs. Since our replication origin detection method (mORI) benefited from higher mutation numbers we conducted it in two stages. In the first stage we identified replication origins based on the PMA score calculated using only the reported TCT→TAT and TCG→TTG variants – Fig.1B.

We then used 1000 identified ORI positions with the highest PMA score to study the distribution around these ORI of all 96 possible triple context-dependent variants. Those that showed similar patterns, belonging to clusters A and B shown on Fig. 2, were used in the second stage of the detection, which was based this time on 58 mutation types. Mutations in cluster A show a positive PMA score while those in cluster B a negative one, in order to account for this difference, in the second stage of the detection we combined mutations with C/A reference allele from cluster A with G/T from cluster B and G/T from cluster A with C/A from cluster B. This increased the total number of mutations used from on average 422k to 606k per sample, also increasing the number of identified replication origins from 5132 to 5409.

**Fig.2:**
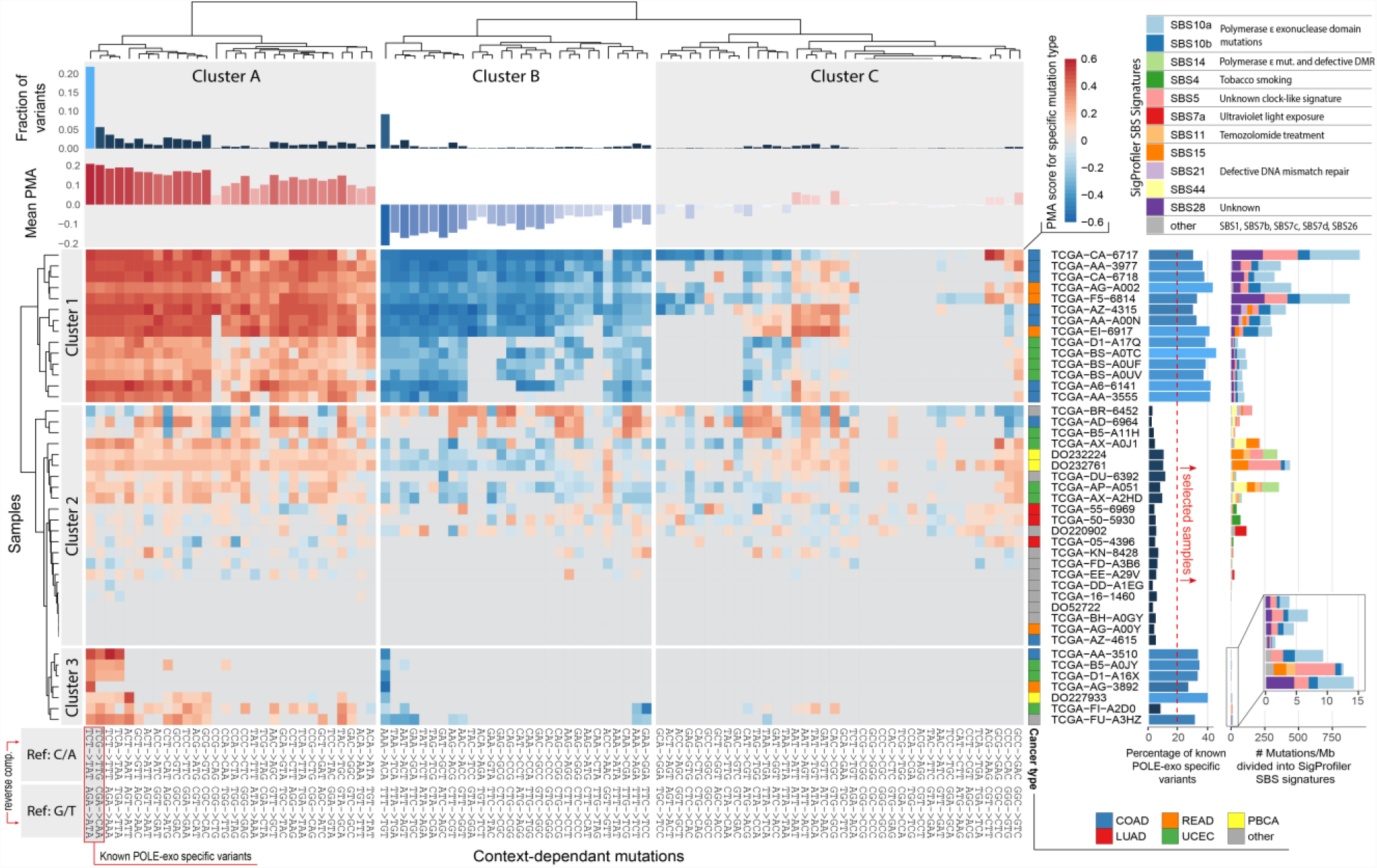
Heatmap of the PMA score calculated at ORI positions identified using known POLE-exo specific variants (marked on the plot). Samples (rows) and mutation types (columns), both clustered into 3 groups. Columns were clustered using only samples in which percentage of POLE-exo specific variants exceeded 20% (bar plot located on the right side). Mutations from clusters A and B were used to calculate PMA score in the final detection. Bar plots on the right show the percentage of POLE-exo specific variants (marked in the lower left corner) and total number of mutations per Mb in each of the samples divided into SigProfiler signatures listed in the upper right corner of the plot. Color-based annotation next to the samples IDs determines the cancer type (COAD - colon adenocarcinoma; LUAD - lung adenocarcinoma; READ - rectum adenocarcinoma; UCEC - uterine corpus endometrial carcinoma; PBCA - pediatric brain cancer).

Fig. 2 shows the PMA score obtained for the 1000 replication origins identified using TCT→TAT and TCG→TTG (referred to as POLE-exo specific variants) for individual mutation types and samples. The majority of 20 selected samples, which show at least 20% POLE-exo specific variants also show a similar pattern in other mutation types, including AAA→ACA / TTT→TGT, which make up around 9% of all mutations in all of the studied samples. Clusters A and B marked on Fig.2 contain a total of 58 mutation types which show different frequency in relation to the replication origin location. Cluster A contains, among others, all C→T mutations no matter the context while cluster B all A→C mutations. The context however seems to affect the frequency of the mutation occurrence with TxT context being associated with the highest number of variants.

The samples differ significantly in terms of mutation profiles. Samples from cluster 1 (Fig.2) show a high number of mutations per Mb, from 52 in TCGA-D1-A17Q up to 952 in TCGA-CA-6717. This cluster represents colon, lung and rectum adenocarcinomas. SigProfiler decomposition of those variants shows a combination of SBS10a, SBS10b, SBS5 and SBS28 signatures the first two of which are characteristic of POLE-exo mutants, the second two are often observed in cancers, especially the SBS5 clocklike signature. Cluster 3 shows a similar SigProfiler decomposition, however the variant frequency is much lower, from an average 2 per Mb in TCGA−AG−3892 up to 14 in TCGA−FU−A3HZ. Cluster 3 contains similar cancer types as cluster 1, with an addition of one cervical squamous cell carcinoma and endocervical adenocarcinoma (TCGA-FU-A3HZ) and one pediatric brain cancer (DO227933). Sample TCGA−FI−A2D0 is an outlier in cluster 3 in terms of mutational signature, showing a majority of variants associated with SBS5 (unknown clock-like signature) and some from SBS15 (defective DNA mismatch repair) unseen in other samples from this group. This sample was not selected for the ORI detection as it doesn’t pass the 20% cutoff of POLE-exo specific variants. Cluster 2 contains the remaining samples which also have less than 20% of POLE-exo specific variants, representing 12 different cancer types. While some of the cases exceed 100 mutations/Mb they show a significantly different pattern associated with defective DNA mismatch repair (signatures SBS15 and SBS44) but also signature associated with polymerase ε mutation (SBS14).

The ORI detection algorithm described is expected to identify only the positions which are conserved among individual cell division cycles and among individuals, except for cell type-specific sites and sites that can change very often depending on the number of cell divisions. We believe these sites are highly dependent on the specificity of the DNA structure in their vicinity, guiding the ORC to those locations.

### Distribution of identified replication origins

Fig.3 shows ORI sites identified in a 3 Mb segment on the p-arm of Chromosome 1, demonstrating the extreme strand bias of selected mutations in their vicinity (the entire list of identified replication origins is available in the Supplementary Table S2). The coefficient also shows local minima, which we assume represent sites where polymerase ε meets polymerase δ, i.e., the endpoints of the replication process. The minima are not as evident as maxima, showing a different distribution and in many cases are not placed halfway between the two neighboring replication origins. Possibly, this could be caused by variation in the firing time of individual replication origins (33), or the polymerase efficiency which, among other things, is known to be GC content-dependent (34).

**Fig.3:**
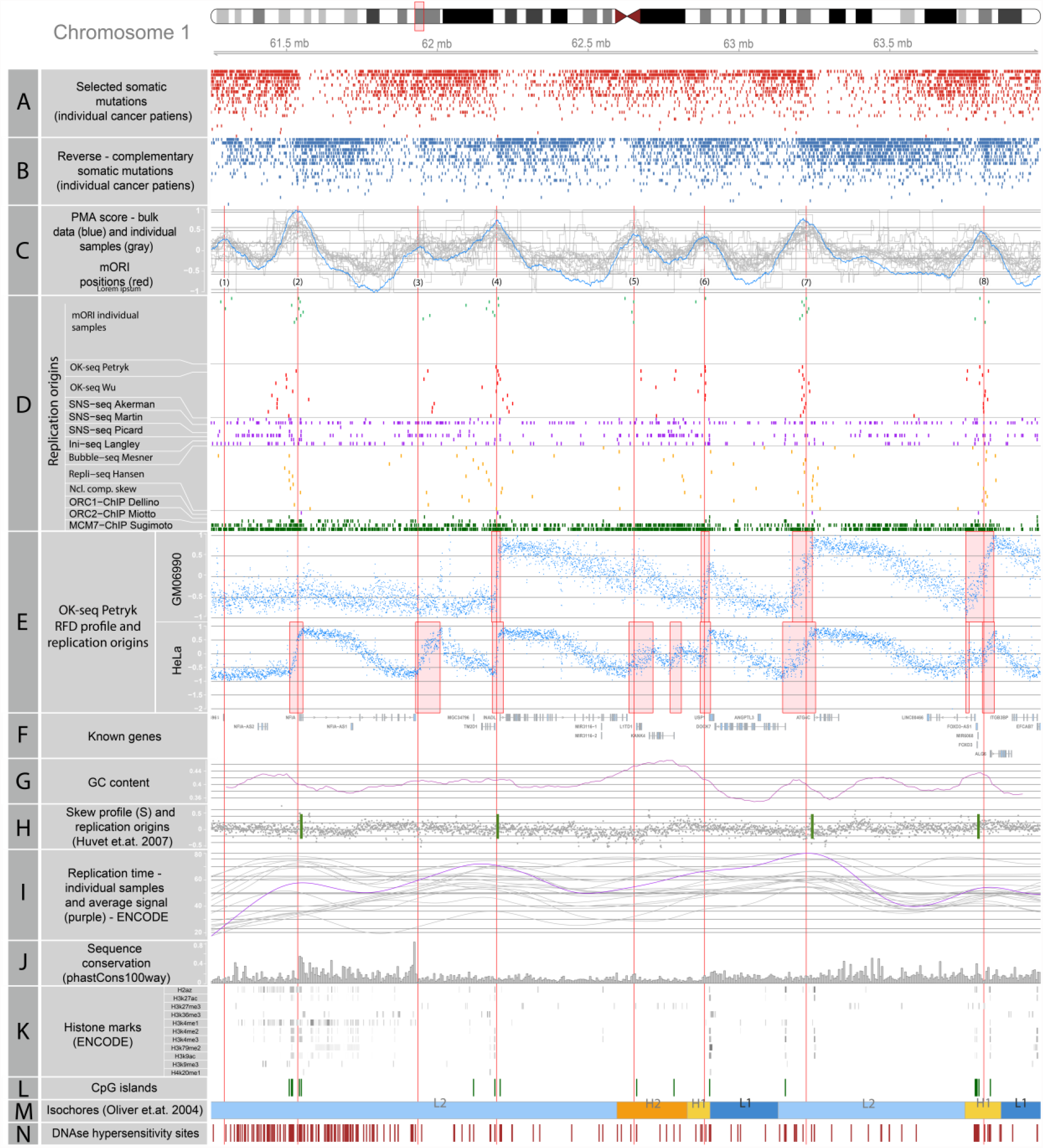
Replication origins identified in a fragment of chromosome 1. **A)** Individual context dependent somatic mutations used in the ORI detection algorithm (C/A reference allele from cluster A and G/T from cluster B, as shown on Fig.2), each row represents one individual patient; **B)** mutations of reverse complementary type to those shown on panel A (G/T reference allele from cluster A and C/A from cluster B as shown on Fig.2); **C)** The consensus PMA score calculated for the combined samples (blue line) and individual sample PMA score (gray), red lines mark the peak positions which represent eight replication origins identified by mORI numbered 1-8; **D)** Replication origins identified by other NGS-based methods, (see Supp Table 1) in other samples from various tissues; each row represents one sample; except the Akerman SNS-seq track where the top one corresponds to core and bottom to stochastic origins **E)** RFD profile (blue dots) for two OK-seq samples from (1), along with identified ORI positions (red rectangles); **F)** exons of known genes; **G)** GC content calculated in 1kb windows; **H)** nucleotide compositional skew profile (gray dots) and replication origin positions from (2); **I)** replication time obtained for individual ENCODE samples (gray line) and an average for all samples (purple line); **J)** sequence conservation score (phastCons100way); **K)** ENCODE histone marks; **L)** CpG islands (UCSC data); **M)** isochore positions from (6), lowest GC - L1,L2,H1,H2,H3 – highest GC; **N)** DNAse hypersensitivity sites obtained for K562 cell line (ENCODE data)

Locations of the replication origins are reported to be associated with DNA features that are non-randomly distributed over the entire genome (7). For this reason, the identified replication origins might also be non-randomly distributed. To test this, we computed the average distance between identified ORI and compared it to the average distance between random positions in the genome. The average distances between the locations in the two sets were then compared, showing a statistically significant difference (*p*-value < 2.2 × 10^−16^), indicating that the variance of the calculated distances is significantly smaller for the identified ORI compared to the random positions, consistent with the former being more regularly spaced.

### Comparison of ORI detection methods

Replication origin detection can be carried out using various experimental approaches, summarized in Table 1, which provide from 1 thousand to over 0.5 million positions in the human genome. The detection is expected to vary significantly in terms of sensitivity and specificity also targeting various classes of replication origins (constitutive, flexible and dormant), resulting in significantly different numbers of detected sites. Our replication origin detection method is based on an asymmetric mutational pattern between the leading and lagging strand. In that respect it is similar to replication fork directionality (RFD) profiles, shown on Fig.3, which originate from the Okazaki fragment purification and sequencing (OK-seq) (1, 35). We compared the mutational pattern in the present work to the experimental RFD profiles available for GM06990, a “normal” cell line, and for HeLa, a cell line of cancerous origin, we also calculated the RFD profiles and carried out origin detection for 10 additional cell lines based on data provided by Wu (35). The number of ORIs detected using our method (5409) is similar to the number described by Petryk et al. for GM06990 (5684) and samples from Wu et al. (on average 5374).

**Table 1.**
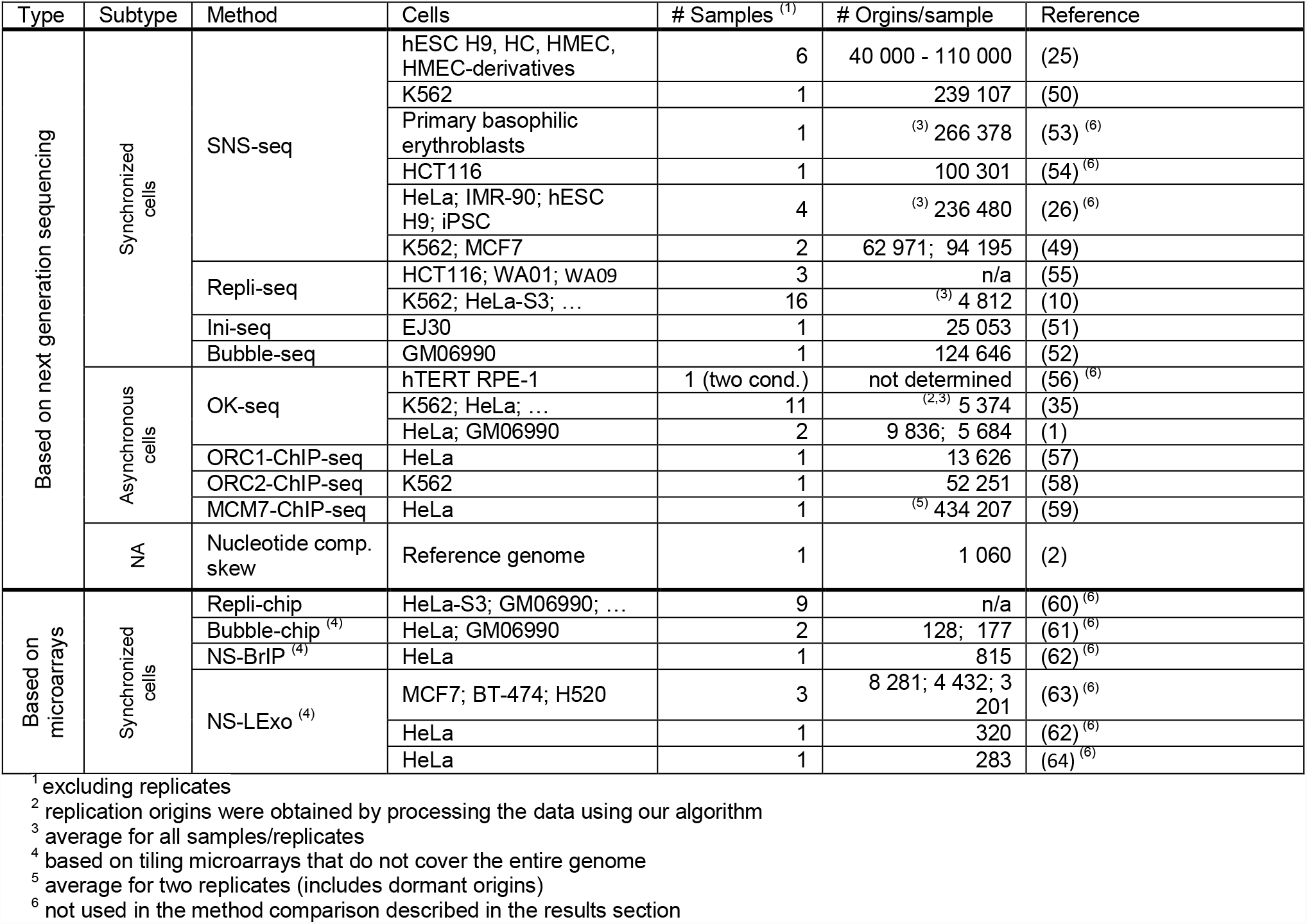
Summary of large scale replication origin detection methods in human cells.

**Table 2:** Sources of alternative ORI positions used in this study

| Method | Reference | Comments |
| --- | --- | --- |
| OK-seq | (35) | BAM files for hg19 were downloaded from the SRA database, under accession number PRJEB25180. Replication origins were identified using our algorithm described below. |
|  | (1) | BED files with ORI positions for hg19 were provided by the authors of the publication. |
| SNS-seq | (25) | Genomic coordinates of ORI positions for hg38 were downloaded from the manuscript supplement - Table S1. The coordinates were converted to hg19 using liftOver function from the *rtracklayer* Bioconductor library (65). ORI from Q1 and Q2 were combined into the "core" group, while the remaining Q3-10 into "stochastic". |
|  | (50) | BED files with ORI positions for hg19, for HeLa, IMR90 and K562 cell lines, were downloaded from http://pbil.univ-lyon1.fr/members/fpicard/oriseq/ |
|  | (49) | hg19 ORI positions were downloaded from the DeOri database (11) for both HCT116 and K562 cell lines |
| Repli-seq | (10) | ENCODE Repli-seq data were downloaded from GEO dataset GSE34399. We used the peak locations of the wavelet-smoothed signal available as the BED files with Pk suffix. |
| Ini-seq | (51) | Genomic coordinates of ORI positions for hg19 were downloaded from the manuscript supplement - Table S3 (Determination by SICER at E = 10e-5). |
| Bubble-seq | (52) | ORI positions for hg18 were downloaded in a bedGraph format from GEO dataset GSE38809 (GM_combined_RD_bubbles file). The coordinates were converted to hg19 using liftOver function from the *rtracklayer* Bioconductor library (65). |
| ORC1-ChIP-seq | (57) | ORI positions for hg18 were downloaded in a BED format from GEO dataset GSM922790 (ChIPseq_Orc1_GradientHela_enrichRegions file). The coordinates were converted to hg19 using liftOver function from the *rtracklayer* Bioconductor library (65). |
| ORC2-ChIP-seq | (58) | Genomic coordinates for hg19 were downloaded from the manuscript supplement (Table S1). |
| MCM7-ChIP-seq | (59) | Genomic coordinates for hg19 were downloaded from GEO dataset GSE107248 (MCM7ChIP_exp1/2.txt files). |
| Nucleotide comp. skew | (2) | ORI positions for hg17 were downloaded from the manuscript supplement (N-domain_positions.xls file). The coordinates were converted to hg19 using liftOver function from the *rtracklayer* Bioconductor library (65). |

**Table 3:**
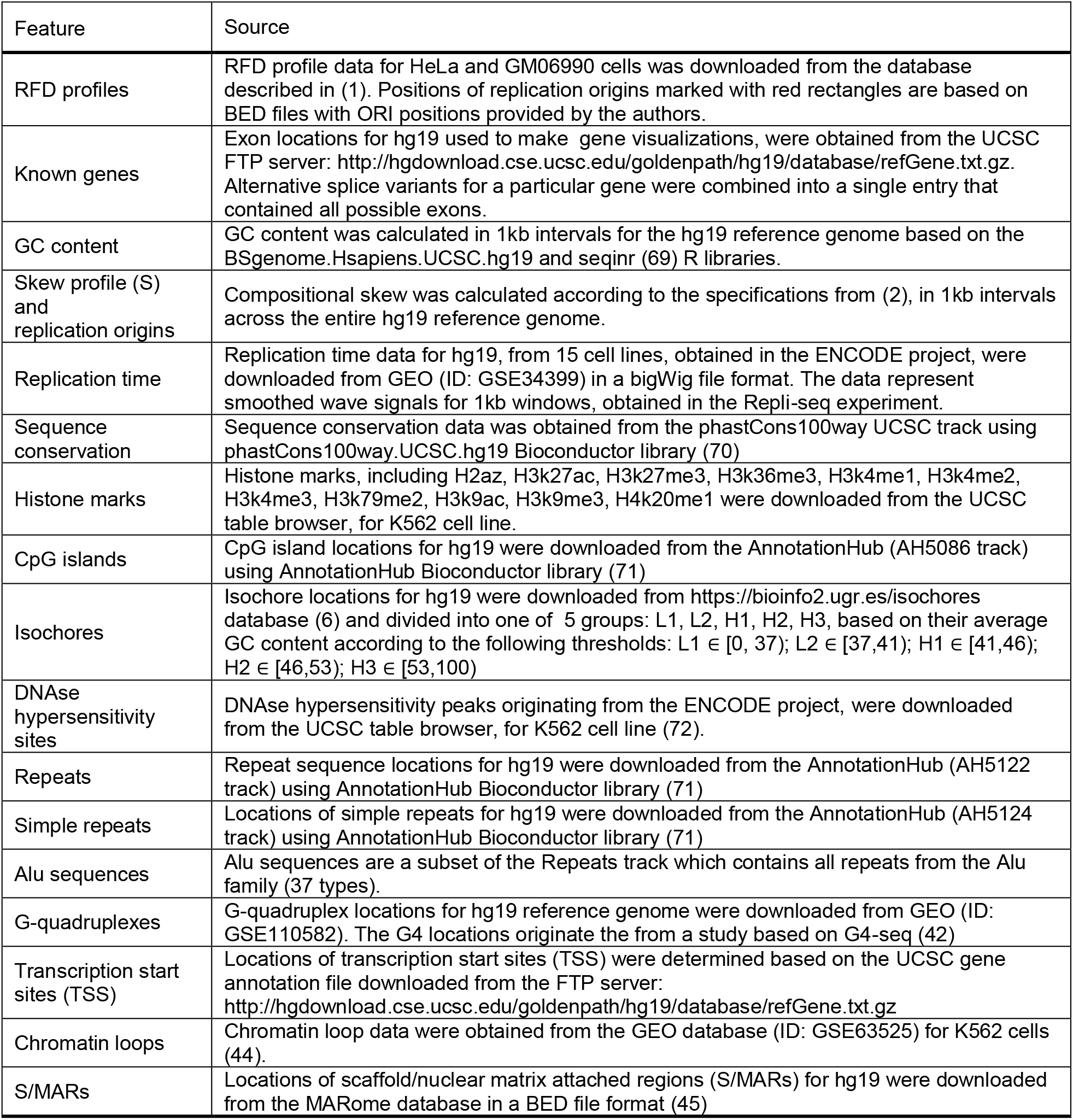
Sources of coordinate locations of various DNA features used in this study.

Comparison of two sets of genomic coordinates is usually carried out by calculating the distance between elements located on a specific chromosome or finding the overlaps after converting the coordinates into ranges (36, 37). This however requires a definition of maximum allowed distance, used to find the overlaps, or pairing the coordinates between both sets for which the distance will be calculated. The latter can be very difficult especially if the overall number of features in both compared sets is different. The problem of comparing two sets of ORI is very similar to the one solved by Needleman-Wunsch global sequence alignment algorithm, where it is possible to introduce gaps into one of the compared sequences in order to account for insertions and deletions. We adopted this dynamic programming approach to work with ORI positions represented as numeric vectors on each chromosome instead of nucleotide sequences and used it to optimally pair to sets of origin positions, which led to an estimation of the pairwise distance between the two sets. Instead of rewards for match and mismatch we used the absolute distance between locations and used a gap penalty to control the number of gaps introduced to optimize the comparison -- gaps represent unmatched positions, from one method or the other (see Materials and Methods). We extended the algorithm to allow comparison of multiple ORI sets, which works similarly to ClustalW multiple sequence alignment (38).

We used both algorithms, termed numeric vector alignment (NVA) and multiple numeric vector alignment (MNVA), to compare the positions of replication origins between methods that provide similar number of replication origins (up to 10 000 positions in the genome). Panels A and B on Fig.4 show the results of MNVA conducted on replication origins we identified, based on mutation patterns in individual samples, with the highest number of mutations, as well as positions detected using the union of all samples. We also used experimentally identified ORI sites that originate from the works of Mesner, Picard, Langley, Petryk and Wu and from the computational method developed by Huvet (see Table 1). Each row represents one origin and the color scale corresponds to the fraction of methods in which it was detected. By applying the MNVA algorithm we were able to collapse nearly 150 thousand genomic coordinates, across all chromosomes, from 31 distinct sets, into 10161 locations. Out of those 4666 were identified in at least 50% and 518 by more than 90% of the samples, from all ORI detection methods. Positions of all compared origins combined using the MNVA algorithm are available as the supplementary table S3. Fig. 4C shows the results of NVA pairwise comparisons between all origins identified. Fig.4 not only shows which positions are identified by the majority or minority of methods (panels A and B) but also the global similarities between them (panel C). Origins detected by mORI were most similar to those detected by the method of Huvet et al. and to the OK-seq. As expected, the three sets of randomly generated genomic positions were poor matches to all other sets.

**Fig.4:**
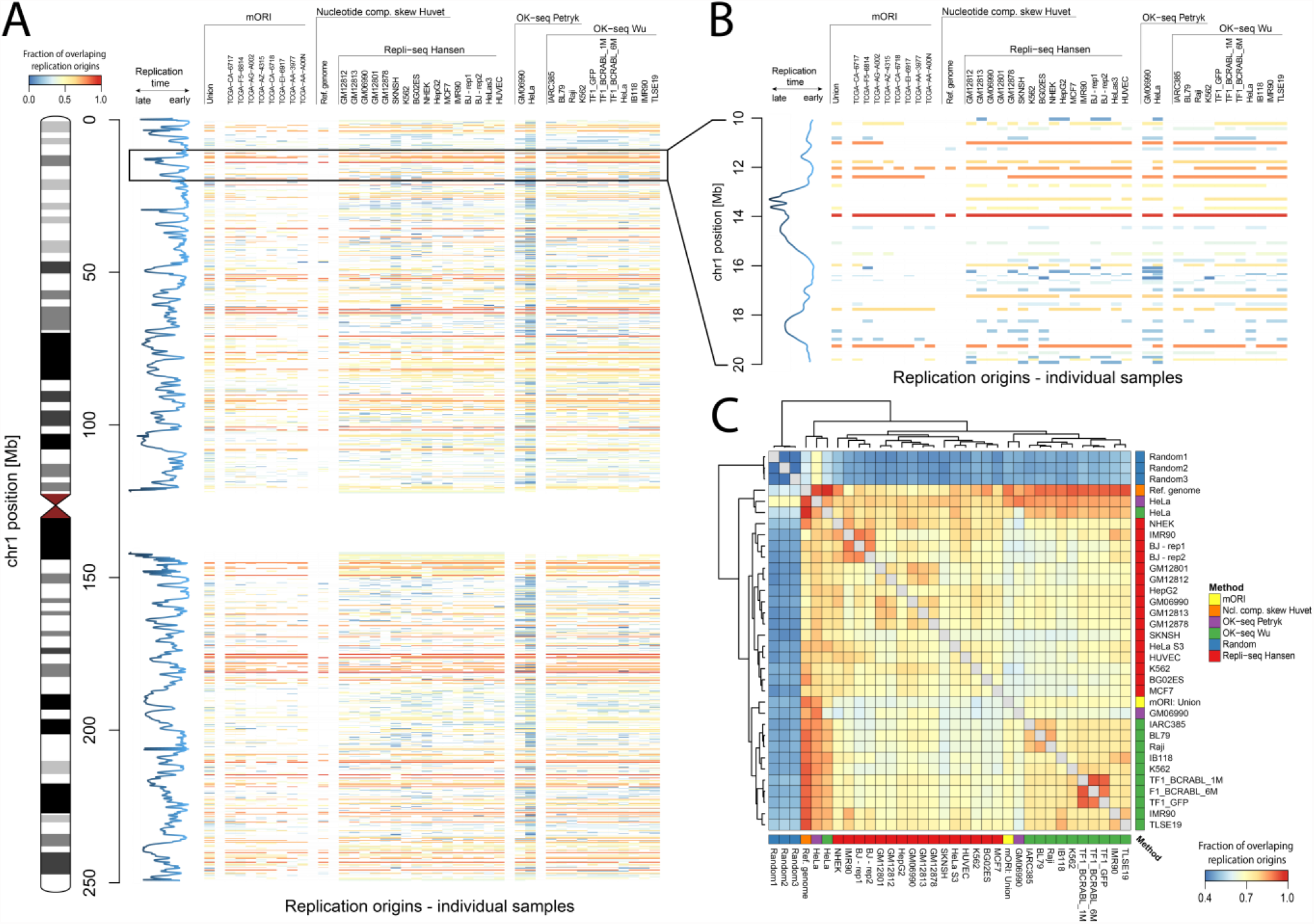
Similarity between various replication origin detection methods. **A**) results obtained using the multiple vector alignment algorithm for chromosome 1, each row represents one origin and the color scale corresponds to the fraction of methods in which it was detected; blue line represents the replication time data. **B**) enlarged fragment of panel A for 10 – 20Mb region of chr1. **C**) heatmap showing overall correlation between ORI detection methods, including a set of 5000 random genomic positions for comparison. The correlations are obtained using a dynamic programming-based vector alignment algorithm (see Methods).

### DNA features associated with ORI

The mORI detection method is based on the PMA score, which additionally can be used to rank the origins. We assume higher PMA scores corresponds not only to the detection confidence but also the frequency at which the origin is utilized in a group of multiple analyzed samples. This provides the possibility to identify a set of consistently utilized replication origin positions that can be used to study the DNA features associated with their location. We conducted such analysis using all 5409 ORI positions, identified based on mutation patterns, and only using a subset of 1000 origins, with the highest values of the PMA score. In Fig. 5A we compared the number of repeat sequences, G-quadruplexes, histone marks, CpG islands, transcription start sites, S/MARs and chromatin loops at various distances from the replication origins, measured by 20kb windows from the putative ORI. The occurrences were summed over all studied origins and divided by the minimum occurrence levels, showing only the fold change for each interval.

**Fig.5:**
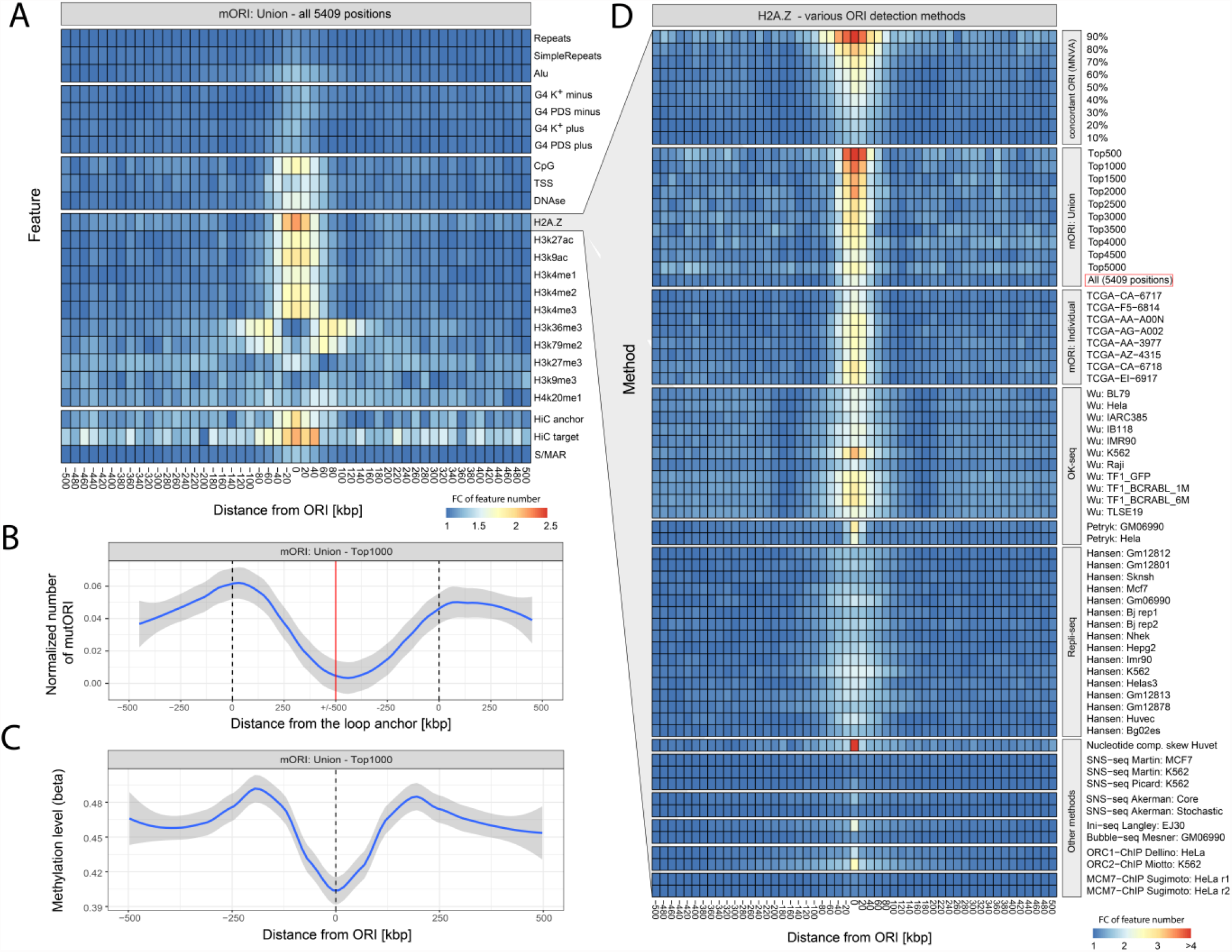
Correlation of mORI sites with epigenetic features of the genome. **A)** Number of specific features at a given distance from the replication origins, expressed as the fold change with respect to the minimum value from each row. **B)** Number of identified ORI at a specific distance from the chromatin loop anchor. Dashed vertical line shows the loop anchor points, red line marks the loop center. Since loops have different size but the plot shows a fixed interval of 1MB, the number of ORI was divided by the number of loops which reach this specific length. **C)** Average methylation level of the TCGA samples used for the detection of replication origins at a various distance from the Top 1000 origin positions. **D)** Statistics for H2A.Z histone obtained for various replication origin detection methods and results of the method concordance obtained using the MNVA algorithm

As expected, the location of replication origins is associated with DNA features that affect the accessibility of DNA to protein binding. Many replication origins are located within promoter and gene-rich regions (7), especially those which are active in a given cell (39). Fig. 5A shows an increase in the number of transcription start sites (TSS) and DNase hypersensitive sites in the vicinity of replication origins (+/-50kb). Stronger association can be observed for the CpG island locations which are also overrepresented in the vicinity or replication origins.

Simple tandem repeats and repeated sequences derived from the Repbase repository (40) show only a minor increase in the vicinity or ORI. The highest increase among repeated sequences is observable for the Alu SINEs, which were reported to be associated with ORI location (10, 41).

G-quadruplexes (G4) were previously reported as being overrepresented in the vicinity of replication origins (26). To test this association we used G4 positions, identified in the (42) study based on G4-seq, which form under physiological K+ conditions and which are stabilized by pyridostatin (PDS). Results provided by both approaches are shown separately for each strand on Fig. 5A. Compared to other approaches, the increase in the G4 number in the vicinity of ORI is small, but clearly observable and offset from center in opposite direction on plus and minus strands.

Promoters of transcriptionally active genes are located inside nucleosome-free regions, which are reported to be associated with replication origins (20, 21). Chromatin structure limits DNA availability, and therefore it is one of the major factors that affect replication origin activity. Among all of the features that we compared H2A.Z histone exhibited the highest occurrence at the replication origins detected by mORI. Long has recently shown H2A.Z histone to epigenetically regulate the licensing and activation of early replication origins (43). Interestingly, our data show that H3k36me3 and H3k79me2 exhibit depletion at the mORI sites, while showing an increased occurrence frequency at the +/- 50-100kb distance.

Topological organization of the DNA inside the nucleus is another important feature that affects DNA replication. Origins located near the anchor points of chromatin loops were shown to have a higher activity (27). To validate this feature we compared the locations of chromatin loops (44), with the locations of the highest scoring 1000 mORI replication origins (Fig. 5B, see Materials and Methods for details). The number of replication origins shows a clear enrichment in the vicinity of loop anchors marked with a vertical dashed line; however, the number of origins drops significantly, approaching the loop center. We also found week association between scaffold/nuclear matrix attached regions (S/MARs), gathered in the MARome database (45), and replication origins which were previously reported to have possible association (28).

The number of mutations associated with the POLE-exo damage is assumed to be correlated with the number of tumor cell divisions (approximately 600 mutations per cell cycle (46)), implying tumor age contributes to the inter-patient differences that were observed (Fig. 3). However, epigenetic modifications to the DNA may also have a significant impact on the location of the replication origins (e.g., DNA methylation (47)) affecting the patient specificity of mutation patterns. We compared the methylation levels, determined in the TCGA project, in the vicinity of mORI replication origins with those for the distant parts of the genome (Fig. 5C). We observed a small drop in the methylation levels in the vicinity of replication origins (∼10%) suggesting that the relation between both is not direct and might result from correlation with other structural features of the DNA.

The PMA score allowed us to select ORI conserved across multiple samples. Using the sample comparison based on MNVA, we also selected a subset of origin positions detected using other approaches. Fig. 5C shows the frequency of H2A.Z histone in the vicinity of replication origins identified using various approaches. The first panel was created using a subset of positions obtained using the MNVA algorithm, based on a various cutoff level, that defines the minimal agreement between all 31 analyzed samples shown below. We excluded the results of our mORI method obtained for individual samples (shown on the “mORI: individual” panel) since the data used to derive them were also used in the combined sample study (mORI: union). The MNVA panel shows that the higher the agreement between the methods, the stronger the association between the replication origin position and frequency of H2A.Z occurrence. A similar trend can be observed in the second panel obtained for the results of mORI selection based on various cutoff levels associated with the PMA score. The higher the PMA score of the identified replication origins, the more frequently H2A.Z is associated with an ORI. While both approaches, based on method concordance and PMA score, can be successfully used to identify a set of conserved origins, the H2A.Z frequency is increased in a narrower interval in the vicinity of ORI selected based on the PMA score and the same situation can be observed for other studied DNA features. This suggests that the origin position is more precisely estimated using the mutation based approach, compared to the agreement between multiple methods, which is why we selected it for the study shown on Fig. 5A-C. Subsequent panels of Fig. 5D show the H2A.Z frequency in the vicinity of origins identified using mORI method applied to individual samples with high mutation counts and other replication origin detection approaches also shown on Fig. 4. Similarly to the ORI identified based on mutation patterns the positions obtained in the OK-seq methods also show high association with H2A.Z, however out of all approaches the most clear association was obtained for the computational method developed by Huvet et al.

## Discussion

We employed on genomic scale the mutator phenotype associated with damaged polymerase ε, found in cancer cases (mainly colon adenocarcinoma and endometrial carcinoma), to identify genomic positions of ORI and developed an approach to compare positions obtained by other means, such as Repli-Seq or OK-seq. Mutations in the exonuclease domain of Polymerase ε affect the proofreading mechanism, leading to characteristic context-specific mutations grouped asymmetrically on the leading strand near ORI. Although exonuclease mutations in POLD1 are much less frequent than in POLE, several cases have been reported (48) and it would be interesting to determine whether POLD1 can also effectively mark ORI.

There are important biological differences in ORI determined by POLE mutational processes compared to approaches depending on isolation and sequencing of newly synthesized DNA, which may lead to deeper insights into the function of mammalian ORI. Mutationally-defined ORI (mORI) are inscribed directly onto the DNA through the action of the replicative enzyme over many rounds of cell division in vivo; Repli/OK-Seq obtain data from a single round of replication in cell culture. One outcome of our mutational approach is that only the most consistently firing origins will be clearly observed, which may reduce the overall ORI count relative to other methods. Repli/OK-Seq are dependent on mapping small fragments of DNA, which may prove limiting in understanding how highly repetitive regions of the genome are replicated. mORI in principle could use long read technologies to overcome mapping difficulties.

We formalized the detection of mutationally-derived ORI by constructing, the POLE-exo mutation asymmetry (PMA) score, which assumes maximum value at the inferred position of ORI. ORI predicted by PMA scores are consistent overall with those obtained by other sequence-based methods. Using a subset of methods that provide a comparable number of replication origins we conducted a two-stage comparison of replication origin positions. In the first stage we conducted a pairwise comparison between each two methods using a numerical vector alignment (NVA) algorithm based on dynamic programming similar to Needleman-Wunsch for global nucleotide sequence alignment. mORI locations were most similar to the OK-seq-derived ORI among sequencing-based method. Subsequently, we compared multiple methods by extending the NVA algorithm to a multiple vector version (MNVA), using an algorithm similar to ClustalW multiple sequence alignment (38). Based on the results we assessed the overlap among all ORI sets and selected origins which are identified in a specific fraction of samples by various methods. We found that 4666 ORI were identified in at least 50% and 518 by more than 90% of the samples, across all methods.

Given the large concordance observed between origins detected using cell types from different patients, we assume that despite cell type and tissue of origin, the majority of ORI positions that we identified can be also observed in normal human cells. We believe that the replication origin positions identified in our study, both based on the mutation patterns and multi-method overlap provide a good basis to study the DNA features that characterize replication origin positions.

Therefore we sought associations of mORI locations with various DNA properties previously shown to co-occur with ORI location. Repeated sequences and G-quadruplexes were weakly associated with mORI origins, while DNA methylation levels and DNase hypersensitive sites were moderately associated. Replication origins from mORI were demonstrated to occur in the vicinity of chromatin loop anchors as was suggested previously (27). Weak association with S/MARs (28) as a potential factor influencing ORI location was also observed. Replication origins identified using mORI were also associated with epigenetic modifications, including methylation level and specific chromatin histones, most importantly H2A.Z. We further used H2A.Z as a benchmark to compare other ORI detection methods as well as various criteria used to select a subset of mORI and ORI identified in multiple samples/methods. Association between ORI and H2A.Z is stronger the higher is our prediction score (PMA) and also stronger as more methods identify a particular origin. Additionally, among the sequencing-based ORI detection methods we showed that Ok-seq, Ini-seq and ORC2-based ChIP-seq exhibit the highest association with H2A.Z. However an even stronger association with H2A.Z was observed for the computational method based on nucleotide composition skew (2). The association with H2A.Z is much weaker for methods that provide high number of origins, especially those based on SNS-seq (25, 49, 50), Ini-seq (51) and Bubble-seq (52). This suggests that they may represent a more variable class of origins—as mentioned, mORI is biased toward constitutive ORI.

The mORI detection method is applicable to cells characterized by the POLE mutator phenotype, which currently is only associated with cancer. This study used those POLE-exo mutated tumors found in ca. 1% of TCGA patients, primarily colorectal and endometrial cancers, but larger studies have been reported with representative tumors from many organ systems including brain, ovaries, prostate pancreas and lung (48). Moreover, it should be possible to introduce exonuclease-mutated POLE into the nuclear genome of any cell type using gene editing technologies.

## Conclusions

We developed a novel method of replication origin detection, MutORI, based on the mutation patterns of POLE-exo tumors. MutORI identifies replication origins using whole genome sequencing data without any modifications, and then classifies origin utilization based on the value of the PMA score, revealing a set of constitutive replication origins in a single step. We applied this methodology to create the first ORI dataset generated from living tissues, rather than cell culture, and used it to characterize DNA structural features in the vicinity of identified ORI positions. The highest association of ORI was with histone H2A.Z, and the association increases with PMA score. We additionally proposed a new replication origin comparison methodologies, for pairwise ORI comparison, based on Needleman-Wunsch global nucleotide alignment, and multiple ORI sets by adapting a ClustalW multiple sequence alignment approach. These methods enabled a comparative analysis of replication origin positions from different cell types and methods. We report the relevant DNA characteristics associated with the locations of the commonly observed ORI. Selection of ORI positions based on our comparison method allowed us to identify 518 highly conserved ORI, which are detected in over 90% of samples from multiple cell lines and using various detection methods.

## Methods

### Detection of replication origins based on somatic mutations (mORI)

For the purpose of the mutation pattern detection we define a POLE-exo mutation asymmetry score PMA which is calculated for each n-th position of the genome:

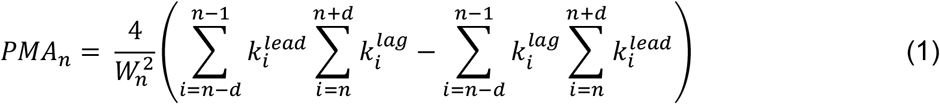

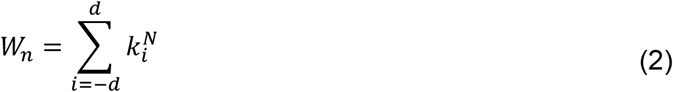

where:

*d* – window size

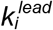 – number of mutations specific to the upstream of replication origin region e.g. T**<u>C</u>**T → T**<u>A</u>**T, identified at position *i* of the genome for all patients

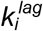 – number of reverse complementary mutations to those from 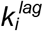 e.g. A**<u>G</u>**A → A**<u>T</u>**A, identified at position *i* of the genome

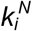 – total number of all mutation types, at position *i* for all patients

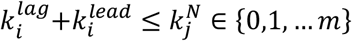

*m –* number of patients

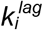 and 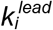 represent mutations specific to the upstream and downstream region from a particular genome position *i* at which the coefficient is calculated. The maximum value of the coefficient is 1 which corresponds to the highest possible difference in mutation occurrence between regions upstream and downstream from a given position. The minimum is −1 which corresponds to a reverse pattern that can be observed in the vicinity of regions where two polymerases meet from opposite directions. The coefficient was calculated every 1kb over the entire genome, using 200kb window (100kb upstream and downstream from the selected position). We then smoothed the coefficient (moving average n=100) and identified local maxima using the peakPick R package (neighlim=100, peak.npos=50). Maxima’s with PMA lower than 0.1 were omitted. The peaks were then filtered using Fisher’s exact test with Benjamini-Hochberg multiple testing correction, assuming 0.01 significance level. This removed regions identified using a small number of mutations which are likely false positives. We tested other approaches to peak filtering including methods utilizing strict criteria based on the PMA score, the number of mutations in the sliding window, or other tests and multiple testing correction methods. All methods used showed a high concordance affecting only the stringency of the tests which is dependent on the parameters and significance levels utilized.

### Replication origin position obtained using other methods

The positions of identified replication origins were compared to positions obtained using other approaches, both experimental and one additional computational. Table 1 summarizes the results showing the number of available samples and identified replication origins. It also highlights which methods were used in the comparison.

Genomic positions of experimentally identified ORI positions were downloaded from 11 various sources, we only used data that originate from NGS based, genome wide studies, listed in Table 1. In some cases the data required some pre-processing, as described in the table below:

### Detection of origins based on OK-seq data

The ORI detection algorithm which we developed for OK-seq data is similar to the approach used in (1). OK-seq is based on replicative incorporation of the EdU followed by size fractionation of EdU labeled fragments and Illumina sequencing. The orgins are identified using RFD profiles, computed for 1kb window using the following formula:

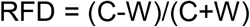

where C and W correspond to the number of reads mapped on Crick and Watson strands respectively. The RFD values range from −1 to 1 which correspond to the highest proportion of leftward and rightward moving forks respectively. In the vicinity of the replication origin the RFD profile should show a significant change in the RFD value from −1 to 1.

To calculate the RFD profile we first used samtools view (66), ver. 1.11, to separate reads from both strands (-b -h -F 16 parameters to get the forward strand reads and -b -h -f 16 for the reverse strand). For this purpose we used aligned reads stored in a BAM file. We than used bedtools coverage (67) ver. 2.25.0 to obtain the number of reads from both strands in 1kb windows defined in a BED file, generated using bedtools makewindows, for hg19 reference genome. Based on those results we calculated the RFD profile using the formula defined above, however we rescaled the number of Crick reads (C), so that the total number of W and C reads is identical for each BAM file. To detect the positions of replication origins based on the RFD profile we first fitted a linear model to the RFD values from a 200kb windows, moving by 1kb across the entire genome. The slope parameter of the linear model has the highest value at positions where the RFD profile shows a shift from −1 to 1, which is assumed to represent replication origins (1). We smoothed (moving average n=200) the slope values of the linear models, calculated for the entire genome and later identified the local maxima, using the peakPick R package (neighlim=100, peak.npos= 100, deriv.lim = 1).

We applied this algorithm to the entire (35) dataset (11 samples) for which positions of replication origins were not provided by the authors. The analysis was based on BAM files downloaded from SRA (PRJEB25180), which were aligned to the hg19 reference genome.

### Comparison of replication origin positions using NVA and MNVA

Genomic coordinates of DNA replication origins were compared using a modified version of the Needleman-Wunch global sequence alignment algorithm, named numeric vector alignment (NVA). The modification allows to use it for comparison of two sets of numeric vectors, instead of nucleotide or amino acid sequences (character vectors). The alignment is performed for each chromosome individually. The modified algorithm doesn’t require a substitution matrix, which defines scores given for matches and miss-matches between specific nucleotides. Instead it uses the absolute distance between both locations and the only parameter required is the gap penalty that controls the number of gaps introduced to the compared vectors.

Individual elements of the scoring matrix *S* are defined as:

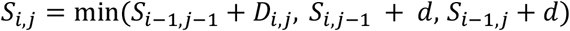

where *D* is the absolute distance between locations *i* and *j* from the A and B vectors:

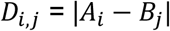

and *d* is the gap penalty parameter. The scoring system is reversed compared to the classical Needleman-Wunch algorithm since the higher is the distance between the elements the less similar are the vectors. The alignment score (value associated with the overall quality of the alignment) is defined as the sum of absolute distances between all paired vector elements. For each unpaired element we add the gap penalty. In our study we used *d*=1000000. The method also returns a consensus vector, which is the average of paired elements from both individual vectors.

To compare a set of multiple replication origin positions, used to create Fig. 4A-B, we created an algorithm similar to the ClustalW multiple sequence alignment (38), named multiple numeric vector alignment (MNVA). The algorithm comprises of the following steps:

1. Calculate all possible pairwise alignments using NVA, record the alignment score for each pair of vectors
2. Create a guide tree based on the pairwise alignment scores matrix obtained in previous step, using the Neighbor Joining algorithm.
3. Align the sequences with NVA by progressive method based on the tree obtained in previous step. In each following iteration the selected vector is aligned to a consensus obtained in the previous iteration.
4. Create the final alignment matrix by aligning each individual vector to the vector obtained in the final iteration of previous step (without introducing new gaps).

The function returns a *n* by *m* matrix where *n* is the number of provided sequences and m is the length of the vector with introduced gaps, obtained in final iteration of step 3. The matrix elements include the original values of each vector separated by added gaps, marked with the dash character. Implementations of both algorithms, NVA and MNVA can be downloaded from our GitHub repository.

### Generation of random genomic positions

Random genomic positions used in the ORI comparisons were selected using the createRandomRegions function from the regioneR package (68), which allows to exclude regions with gaps in the reference genome. For the study of distances between ORI identified using PMA score which was compared to random genomic positions we additionally selected the positions only from regions were mutations were identified. This approach allows omitting the regions with unknown sequence and low complexity, which due to lack of mutation data would bias the results.

### Analysis of genomic features in the vicinity or replication origins

The following table lists all genomic features visualized on Fig. 4 and used to determine the number of features at a specific distance from replication origins used to create Fig. 5A,D:

### Study of methylation levels at ORI locations

Processed methylation data (beta values) for hg19 were downloaded from the GDC Data Portal (TCGA project) for 19 out of 20 samples, which were used for replication origin detection based on mutation pattern. The only omitted sample was DO227933, for which methylation data was not available in ICGC database at the time we wrote the manuscript. The downloaded data originated from both Illumina Infinium HumanMethylation450 and HumanMethylation27 BeadChips. While both platforms differ significantly in the number of tested CpGs we were only interested in the variability of methylation levels at a certain distance of identified replication origins without a direct comparison between the samples. For this reason all samples were combined into one set which was used to calculate average methylation levels at a particular genomic location. The differences between both platforms affected the weight of individual sample in the average methylation levels, however we find that to be of small relevance to the conclusions we have drown based on those statistics.

### Comparison of ORI locations and chromatin loops

Chromatin loop data were obtained from the Gene Expression Omnibus database (ID: GSE63525) for K562 cells (44). The locations were used to estimate the distribution of replication origins in the vicinity of loop anchor by calculating the number of replication origins at a specific distance from the closest loop. We considered the loop itself and the +/- 500kb surrounding region; however, since the loops have different lengths we divided the number of origins by the total number of loops that reach a specific length, similarly as we proposed in our previous work (73) for transcription factor binding sites.

## Abbreviations

G4: G-quadruplexes
ICGC: International Cancer Genome Consortium
MNVA: Multiple numeric vector alignment
mORI: mutationally-defined ORI
NVA: Numeric vector alignment
ORI: origins of replication
PMA: POLE-exo associated Mutation Asymmetry score
RFD: replication fork directionality
S/MARs: scaffold/nuclear matrix attached regions
TADs: topologically associating domains
TCGA: The Cancer Genome Atlas
TSS: transcriptions start sites
WGS: whole genome sequencing data

## Declarations

### Ethics approval and consent to participate

Not applicable

### Consent for publication

Not applicable

### Availability of data and materials

Details concerning POLE-exo mutants, positions of identified replication origins and comparison of replication origin positions, based on MNVA algorithm, identified using multiple approaches are available as supplementary tables.

Implementations of the main R functions used in this study are available in the replicationOrigins R package, which is available through GitHub: https://github.com/rjaksik/replicationOrigins. The package includes implementations of the following algorithms:

- Replication origin detection based on mutation patterns
- OK-seq origin detection based on RFD profiles
- Numeric vector alignment (NVA)
- Multiple numeric vector alignment (MNVA)

The remaining code used for additional analysis and visualizations is available per reasonable request.

### Authors’ contributions

RJ implemented the data processing methods, conducted the analysis and drafted the paper. DW contributed to the study design, interpretation of the results and writing of the manuscript. MK supervised the project and contributed to the study design, aided the data analysis and contributed to the writing of the manuscript.

## Acknowledgements

The results shown here are in part based upon data generated by the TCGA Research Network: https://www.cancer.gov/tcga. Calculations were carried out using infrastructure of the Ziemowit computer cluster (www.ziemowit.hpc.polsl.pl) in the Laboratory of Bioinformatics and Computational Biology, The Biotechnology, Bioengineering and Bioinformatics Centre Silesian BIO-FARMA, created in the POIG.02.01.00-00-166/08 and expanded in the POIG.02.03.01-00-040/13 projects.

